# Anti-MDA5 monoclonal antibodies from patients with dermatomyositis - B cell characteristics and differential targeting of the helicase domains

**DOI:** 10.64898/2025.12.01.691530

**Authors:** Eveline Van Gompel, Marina Galesic, Nicolas Delaroque, Karolin Kern, Ragnhild Stålesen, Antonella Notarnicola, Deniz Demirdal, Monika Hansson, Annika van Vollenhoven, Edvard Wigren, Susanne Gräslund, Michael Szardenings, Vivianne Malmström, Ingrid E Lundberg, Begum Horuluoglu, Caroline Grönwall, Karine Chemin, Vijay Joshua

## Abstract

**Objectives:** Autoantibodies targeting melanoma differentiation associated protein 5 (MDA5) are strongly associated with dermatomyositis (DM) and may contribute to its pathogenesis. Here we aimed to investigate MDA5^+^ B cells, their phenotype and generate MDA5 monoclonal antibodies to assess their epitope specificity.

**Methods:** MDA5-reactive B cells were captured from peripheral blood of patients with anti-MDA5^+^ DM (n=3) using an MDA5-fluorescent probe. B cell receptor (BCR) sequences were analysed from single-sorted B cells (n=240). Selected clones were re-expressed as IgG1 monoclonal antibodies (mAbs, n=23). Reactivity was assessed using recombinant MDA5 protein constructs, peptide epitope mapping, ELISA, western blot and a commercial line blot assay.

**Results:** Of 240 anti-MDA5^+^ sorted B cells, 23 BCRs were re-expressed as mAbs, two of which showed high reactivity and specificity for MDA5. These antibody sequences originated from one CD19^+^IgD^-^CD27^-^CD38^+^ and one CD19^+^IgD^-^CD27^+^CD38^+^ IgG^+^ B cell with low somatic hypermutation (SHM). Both mAbs had nanomolar apparent affinity and bound to sites within the helicase domains of the MDA5 protein but with distinct epitope recognition. Serology screening confirmed targeting of a linear epitope identified in the mAb studies.

**Conclusion:** Our results show that anti-MDA5^+^ B cells recognize the helicase domains, which are the enzymatically active domains of the protein. These results have implications for understanding the etiopathology of anti-MDA5^+^ DM and development of new antigen-specific therapies.

## INTRODUCTION

Dermatomyositis (DM) associated with IgG autoantibodies to melanoma differentiation-associated protein 5 (MDA5) is a rare, systemic autoimmune disorder that mainly affects the skin with characteristic ulcerations and often with muscle weakness, arthralgia/arthritis, and interstitial lung disease (ILD). Importantly, some patients develop severe rapidly progressive interstitial lung disease (RP-ILD) that can result in death due to respiratory failure [1–5]. The MDA5 protein is a highly specialized intracellular pattern recognition receptor that upon binding of (viral) dsRNA polymerizes and induces type I interferon (IFN) genes that will initiate an inflammatory response [1, 6–8]. Anti-MDA5 autoantibody serum levels may fluctuate and potentially increase upon clinical flares and decrease or disappear with improvement of the clinical disease manifestations [9, 10]. The anti-MDA5 autoantibodies can therefore serve as a biomarker for both disease activity and severity [11, 12]. These observations provide indirect evidence for a role for the anti-MDA5 autoantibodies in the disease pathogenesis, but the underlying molecular mechanisms remain to be understood. Here we identify and phenotype MDA5^+^ B cells, and clone their receptors to produce monoclonal antibodies, with the aim to better understand their targeted epitope and autoreactivity in DM.

We utilized flow cytometry to detect and phenotype MDA5-reactive B cells in the peripheral blood of patients with MDA5^+^ DM. Next, single autoreactive B cells were sorted and subjected to B cell receptor (BCR) sequencing, and patient-derived anti-MDA5 monoclonal antibodies were generated and characterized. The MDA5-reactive mAbs originated from antigen-experienced B cells and identified new linear epitopes within the helicase 2i (Hel2i) domain targeted in patients with DM.

## METHODS

### Patients and Samples

Peripheral blood mononuclear cells (PBMC) for cell studies (n=3) and serum samples (n=21) for serology, from anti-MDA5 positive patients diagnosed between 1999 and 2021 at Karolinska University Hospital (Stockholm, Sweden) were used. Patients were classified as DM according to the EULAR/ACR criteria [13] and found to be anti-MDA5^+^ by a commercially available line blot assay (Euroimmun), and/or by immunoprecipitation (IP) in combination with enzyme-linked immunosorbent assay (ELISA) [14]. A summary of demographic and clinical findings is available in Figure 1A and Table 1. Physiciańs global disease activity was based on a visual analogue scale (0-100). ILD was defined by radiological findings of inflammation or scarring (fibrosis) of the lung parenchyma by high resolution computerized tomography (HRCT) in combination when available with results from pulmonary function tests as described previously [15]. Creatine kinase (CK), aspartate-amino-transferase/alanine-amino-transferase (ASAT/ ALAT), C reactive protein (CRP) and erythrocyte sedimentation rate (ESR) were all determined at the time of blood sampling.

**Figure 1.**
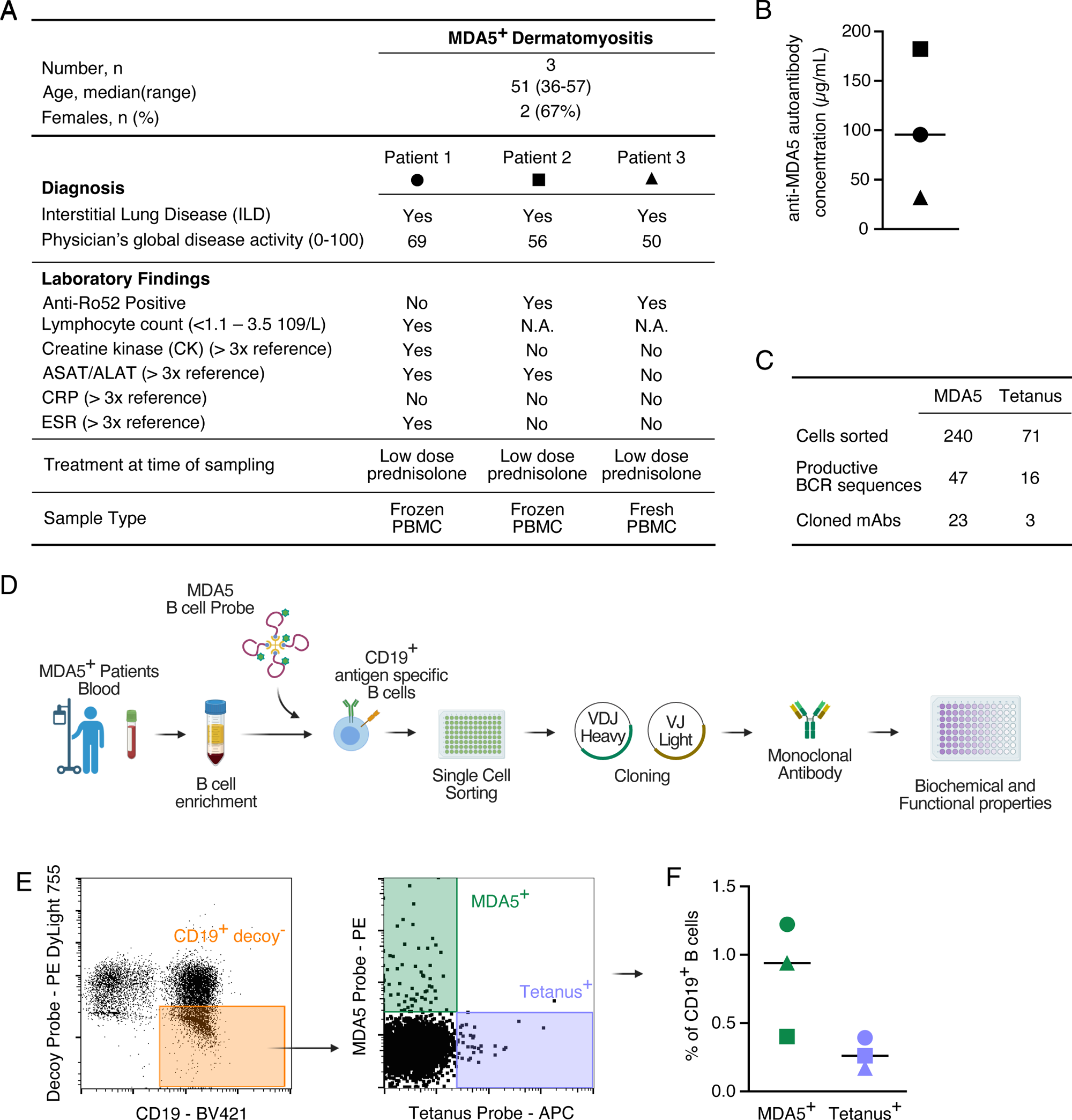
Detection, isolation and phenotyping of anti-MDA^+^ autoreactive B cells. **A and B)** Characteristics and MDA5 antibody levels of the three patients from whom antigen specific B cells were isolated **C)** The table displays the number of sorted cells, the number of complete BCR heavy and light chain sequence pairs and the number of cloned monoclonal antibodies (mAbs). **D)** Schematic overview of the antibody cloning process explained in detail in the supplementary Methods. **E)** Representative flow cytometry plots showing the gating strategy to identify and single sort antigen specific (MDA5 or tetanus) CD19^+^ B cells from PBMC. **F)** Proportion of antigen-specific B cells among the CD19^+^ B cells.

**Table 1:** Demographics of Patients and Healthy individuals.

|  | **MDA5 Positive** | **Healthy** |
| --- | --- | --- |
| n | 21 | 93 |
| Age at diagnosis, median (IQR) | 41 (36-51) | 58 (47-67) |
| Sex (female), n (%) | 8 (38.1) | 58 (62.4) |
| Diagnosis |  |  |
| PM, n (%) | 3 (14.29) | NA |
| DM, n (%) | 16 (76.19) | NA |
| ADM, n (%) | 2 (9.52) | NA |
| Myositis associated autoantibodies |  |  |
| Anti-SSARo52 positive, n (%) | 11 (52.4) | NA |
| Disease course |  |  |
| Disease duration (months), median (IQR) | 2 (1-3) | NA |
| Immunosuppressive treatment, n (%) | 15 (71) | NA |
| Clinical symptoms at diagnosis |  |  |
| Muscular manifestations |  |  |
| Muscle weakness, n (%) | 11 (55) | NA |
| MMT8, median (IQR) | 79 (70-80) | NA |
| Elevated muscle enzymes, n (%) | 12 (66.7) | NA |
| Dermatological manifestations |  |  |
| Heliotrope rash, n (%) | 11 (52.4) | NA |
| Gottron's papules/sign, n (%) | 18 (85.7) | NA |
| Respiratory manifestations |  |  |
| TLC <80%, n (%) | 12 (70.6) | NA |
| FVC <80%, n (%) | 13 (72.2) | NA |
| DLCO <75%, n (%) | 9 (69) | NA |
| ILD, n (%) | 18 (85.7) |  |
| RP-ILD, n (%) | 3 (15) | NA |
| Radiographic lesions, n (%) | 18 (85.7) | NA |

SD, standard deviation; PM, polymyositis; DM, dermatomyositis; ADM, amyopathic dermatomyositis; IQR, interquartile range; TLC, total lung capacity; FVC, forced vital capacity; DLCO, diffusing capacity of the lungs for carbon monoxide; ILD, interstitial lung disease; RP-ILD, rapid progressive interstitial lung disease; NA, not applicable.

Additional serum samples (n=93) were obtained from healthy individuals with no self-reported autoimmune diseases as controls. All individuals and patients who donated samples included in this study provided written informed consent for the use of the samples for research purposes and the ethical permits were issued by the regional ethics committee in Stockholm (Sweden).

### Generation of recombinant MDA5 protein constructs

MDA5 constructs spanning different MDA5 domains were generated as previously described [16]. Biotinylated recombinant proteins were expressed in *E.coli*, purified by a two-step procedure including immobilized metal affinity chromatography (IMAC) followed by size exclusion chromatography (SEC) and evaluated by SDS-PAGE (NuPAGE Bis-Tris 4–12%, Invitrogen). Construct IDs and amino acid coverage are available in Supplementary Table 1. Retinoic acid–inducible protein I (RIG-I) was expressed as a negative control.

### Flow cytometry analysis and capture of MDA5^+^ B cells

Peripheral blood mononuclear cells (PBMC) were isolated using Ficoll-Paque Plus (Cytiva) density gradient centrifugation. B cells were enriched from fresh (n=1) and cryopreserved (n=2) PBMC using a negative selection kit (EasySep™ Human B Cell Enrichment Kit II Without CD43 Depletion, Stemcell Technologies) according to the guidelines of the manufacturer. Fluorescently labelled probes for detection of antigen-specific B cells were generated by loading the biotinylated 88.7 kDa MDA5 protein (covering residues A298-D1025 and domains Hel1-Hel2i-Hel2-pincer-CTD) onto streptavidin-phycoerythrin (SA-PE) (Agilent Technologies) in a 10:1 molar ratio. Free MDA5 was removed by ultra-filtration (Amicon^®^ Ultra, 100 kDa MWCO, Merck). As a control, biotin-tetanus toxoid (TT; Mabtech AB) was coupled to SA-Allophycocyanin (SA-APC; Agilent Technologies), and as decoy, DyLight^TM^ 755 (DL755, Thermo Fisher Scientific) was covalently conjugated to SA-PE.

Enriched B cells were stained at 4°C in three steps: first with Zombie NIR™ live/dead stain (BioLegend Inc.) for 10 min; second with SA-PE-DL755 decoy probe (25 nM) for 20 min; and finally with MDA5-SA-PE and tetanus-SA-APC probes (10 nM) in combination with fluorescently labelled antibodies against cell surface receptors for 20 min (see Supplementary Table 2). Single CD19^+^Decoy^-^MDA5^+^/tetanus^+^ B cells were index-sorted using a BD Influx™ Cell Sorter (BD Bioscience) by manual gating and collected in a 96 well PCR plate.

### BCR sequence analysis and expression of monoclonal antibodies

Heavy and light chain immunoglobulin sequences were obtained from single cells by multiplex PCR as previously described [17]. Sequences were annotated using IMGT/V-QUEST [18]. The similarity between the sequences were analysed by combining the heavy and light chain sequence and performing a multiple sequence alignment [19] and visualised using ggtree (R package) [20]. Estimation of antigen-driven selection pressure for immunoglobulin sequences was obtained using BASELINe tool, as previously described [21]. Variable region antibody sequences of the MDA5-reactive BCRs were de novo synthesized (Integrated DNA Technologies) and cloned into heavy (γ1) and light chain (k/l) antibody expression vectors. Recombinant hIgG1 was expressed in Expi293F cells (Thermo Fisher Scientific), purified by protein G (Cytiva) affinity chromatography, and extensively quality controlled using SDS-PAGE, ELISA and size exclusion chromatography. Non-specific binding was assessed using a polyreactivity ELISA based on HEK293 soluble membrane protein fraction [22], a commercial line blot (Myo4, Euroimmun), and a custom-designed solid phase antigen array (Thermo Fisher Scientific, ImmunoDiagnostics).

### MDA5 Enzyme-Linked Immunosorbent Assay and Western Blot

Binding of mAbs against MDA5 and its different domains was assessed using in-house ELISA and Western blot as previously described [16].

For ELISA, half-area high binding plates were coated with neutravidin 1 μg/ml (Sigma-Aldrich), blocked with 1% BSA PBS, and MDA5 proteins were captured at 0.5 μg/ml. mAbs were analyzed at 5 μg/ml for screening and thereafter titrated for comparison of binding reactivity. Binding was detected with HRP-conjugated goat anti-human IgG F(ab)2 (Jackson ImmunoResearch) and 3,3,5,5-tetramethylbenzidine (TBM) substrate (Sigma-Aldrich).

For Western blot, 250 ng of the different MDA5 protein constructs were separated on 4-12% Bis-Tris NuPAGE with MES-SDS running buffer (ThermoFisher Scientific) and transferred to PVDF membranes. Antibody binding was assessed at 5 μg/ml and detected with HRP-conjugated goat anti-human IgG F(ab)2 (Jackson ImmunoResearch) and SuperSignal^TM^ West Pico chemiluminescent substrate (Thermo Fisher Scientific).

The apparent affinities of anti-MDA5 monoclonal antibodies were estimated by in-solution binding of IgG1 mAb (at a constant concentration of 0.25 μg/ml) and biotin-MDA5 protein (ranging between 228-0.009 nM) for 1hr at room temperature (RT). Complexes were then captured on protein G (Thermo Fisher Scientific) coated ELISA wells for 20 min and detected using streptavidin-HRP (Jackson ImmunoResearch) secondary antibody. EC_50_ or IC_50_ was calculated by interpolating of the optical density (OD) vs concentration on a sigmoidal four parameter logistic curve.

### Epitope-mapping of anti-MDA5 monoclonal antibodies

Epitope mapping of the MDA5 monoclonal antibodies was performed using custom synthesized peptides on cellulose-ßalanine PepSpots membranes (JPT Peptide Technologies). The array consisted of 117 spots containing 5 nmol covalently bound fifteen-mer peptides, overlapped by eleven amino acids covering the C-terminal of MDA5 starting at the Hel2i domain (P549 to D1025, Acc. No. Q9BYX4). Membranes were rinsed in methanol, washed with 0.05% Tween 20 Tris-buffer saline (TBS-T) and blocked (5% BSA TBS-T) overnight at 4°C. mAbs at 5μg/mL were then incubated for 3 hours at RT with agitation. Following washes, membranes were incubated for 2 hours with goat anti-human IgG HRP-conjugated (Jackson ImmunoResearch). Antibodies were diluted in 0.1% BSA supplemented TBS-T. SuperSignal West Pico substrate (Thermo Fisher Scientific) and ChemiDoc Imaging system (Bio-Rad) were used for detection.

Secondly, the clones were subjected to epitope fingerprinting using peptide phage display methodology as previously described [23]. Briefly, two rounds of phage display from a specialized naïve 16-mer peptide library were performed using anti-MDA5 mAbs bound to protein A Dynabeads (Thermo Fisher Scientific). Recovered phagemids were sequenced by Illumina MiSeq next-generation sequencing (NGS). Statistics for 3-mer and 4-mer peptide motifs were calculated, and mapped to the human MDA5 protein sequence to identify peptides sharing potential epitope motifs. Peptide candidates were synthesised with N-terminal biotin and evaluated by ELISA using the same protocol as for the MDA5 protein constructs.

### Peptide ELISA and serology screening of identified binding epitopes

Serum IgG reactivity against the identified novel MDA5-Hel2i peptide epitope was screened in patients with anti-MDA5^+^ dermatomyositis using a cyclic biotinylated peptide MDA5_644-656_ (Biotin-Axh-HQ**C**-DSDEGGDDEY**C**DG) (GenScript) and an in-house ELISA. 96 well half area high-binding plates (Corning) were precoated with NeutrAvidin (Thermo Fisher Scientific) in PBS, blocked with 1% BSA in PBS and bio-MDA5_644-656_ peptide was captured at 0.5 μg/mL in 0.1% BSA-PBS-0.05% Tween 20. Patient and healthy control sera were diluted 1:100 in radioimmunoassay buffer (10 mM Tris-HCl, 1% BSA, 1% Triton X100, 0.1% SDS, pH 7.6) and bound antibodies were detected using goat anti-human IgG HRP (Jackson ImmunoResearch) and TMB substrate (Sigma-Aldrich). OD obtained from serial dilutions of 223:01F12 were used for generating standard curves that allowed for interpolation of anti-MDA5_644-656_ serum values (in μg/mL) using four parameter logistic curve. 98th percentile of healthy cohort (n=93) serum reactivity was set as cut-off value for autoantibody positivity.

### Statistical analyses

The distribution of the data was assessed by performing D’Agostino-Pearson test, Shapiro-Wilk test or Kolmogorov-Smirnov test depending on the sample size. Comparisons between two groups were performed using parametric T-test or non-parametric Mann-Whitney U test, when appropriate. The significance was defined at p value less than 0.05. Data are reported as mean and standard deviation (SD) or median and interquartile range (IQR), when appropriate. GraphPad Prism 9.0 (GraphPad Software) was used for statistical analysis and graphical visualisation.

## RESULTS

### MDA5^+^ B cells were identified in peripheral blood of patients with MDA5^+^ dermatomyositis

Peripheral blood from three patients with MDA5^+^ DM and ILD available from time of diagnosis was analysed (Figure 1A). Serum concentrations of anti-MDA5 autoantibodies were measured by in-house ELISA [16] and ranged from 32-182 µg/mL (Figure 1B). After B cell enrichment, MDA5^+^ and Tetanus^+^ CD19^+^ B cells without decoy reactivity were single cell sorted (Figure 1C, D, E and supplementary Figure 1A). The frequency of MDA5^+^ B cells compared to tetanus^+^ B cells was higher in all patients with DM (Figure 1F). Notably, the MDA5 probe, like the tetanus probe, captured both naïve and memory B cells. The proportion of naïve (IgD^+^ CD27^-^), switched memory (IgD^-^ CD27^+^), unswitched memory (IgD^+^ CD27^+^) or double negative (IgD^-^ CD27^-^) B cells were similar among the two specificities (supplementary Figure 1B). However, it should be acknowledged that the number of analysed anti-tetanus cells was limited. It should also be noted that one of the patients with DM (patient 3) displayed a substantially skewed blood cell profile with high numbers of unswitched memory B cells (supplementary Figure 1C).

From the three patients, we single sorted 240 CD19^+^ MDA5^+^ B cells and 71 CD19^+^ tetanus^+^ B cells (Figure 1C). PCR amplification of BCR transcripts generated 63 anti-MDA5 complete heavy and light chain sequence pairs. Compared to the anti-tetanus repertoire, anti-MDA5+ B cells showed increased usage of *VH* genes such as *VH3-23*, *VH3-74*, *VH4-34*, *VH4-39* and *VH5-51* (supplementary Figure 2A and B). There were two subsets of MDA5^+^ cells with either lower or higher somatic hypermutation (SHM) levels (supplementary Figure 2C).

### Monoclonal MDA5 antibodies were generated from MDA5^+^ patient B cells

From the obtained single cell BCR sequences, 26 clones (23 MDA5^+^ and 3 tetanus^+^) were selected for re-expression as recombinant IgG1 monoclonal antibodies (mAb) (Figure 1C and supplementary Figure 3A). The sequences of the expressed mAbs originated from five IgA^+^, 10 IgG^+^ and 11 IgM^+^ B cells (Figure 2A). Of the 26 expressed mAbs, 12 (46.2%) showed varying degrees of polyreactivity and six (23.1%) showed varying degrees of MDA5 reactivity as assessed by an in-house ELISAs (Figure 2A). Two mAbs 223:01F01 (F01) and 223:01F12 (F12) showed high MDA5 positivity and no polyreactivity and were selected for further studies.

**Figure 2.**
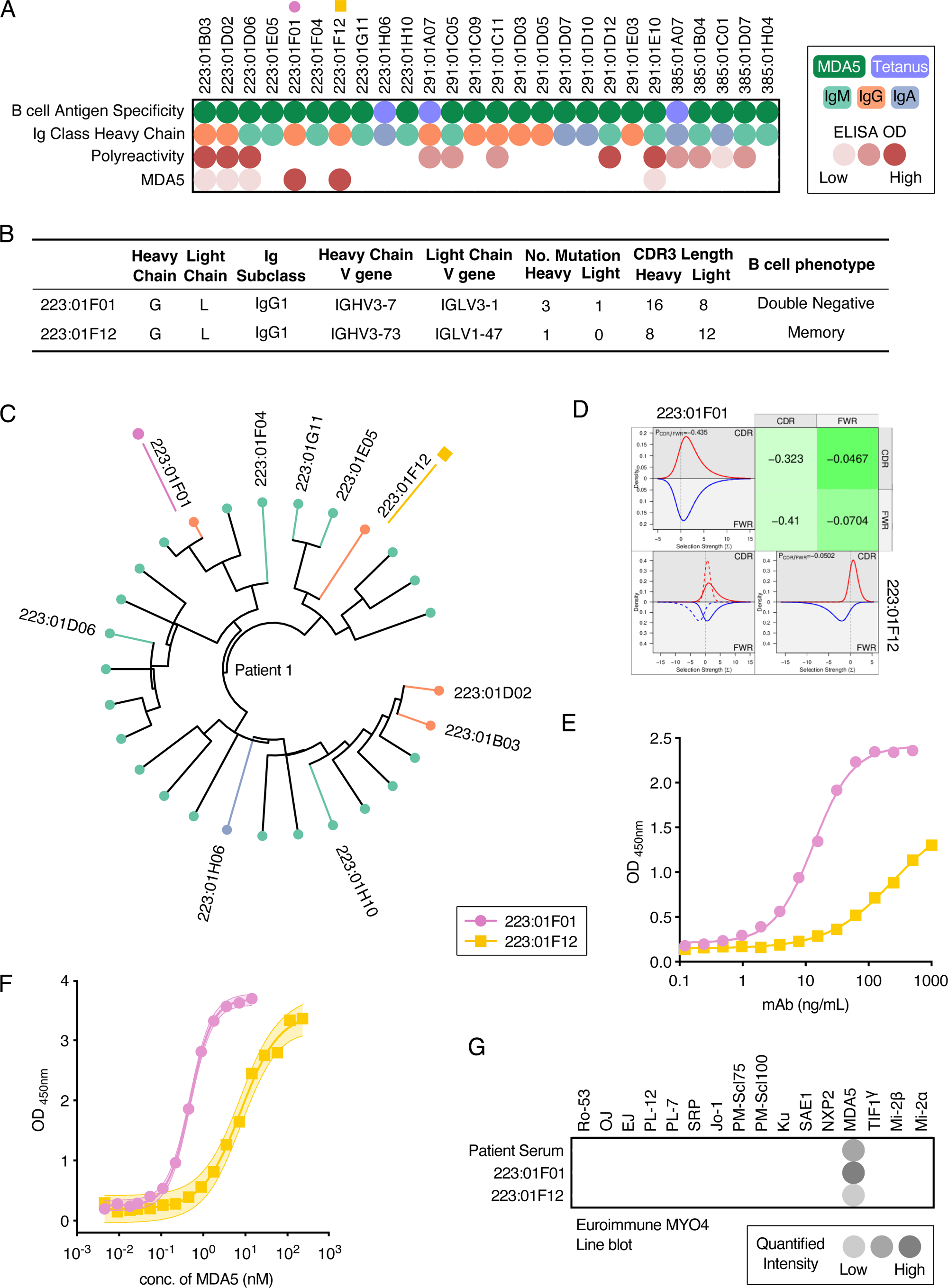
Generation of patient-derived monoclonal antibodies to MDA5. **A)** The heatmap shows the characteristics of the 26 BCRs expressed as mAbs. The map shows the antigen specificity of the sorted B cells (MDA5^+^ or tetanus^+^), Ig isotype (A, G or M) of the BCR, polyreactivity and MDA5 reactivity based on ELISA. **B)** Immunogenetics of the identified MDA5^+^ clones **C)** Phylogenetic tree depicting the relationship between the combined heavy- and light- chain Ig sequences of the sorted antigen specific B cells from a MDA5^+^ dermatomyositis patient in whom the two MDA5^+^ mAbs were identified. Each node represents an individual heavy and light chain. Sequences corresponding to expressed mAb are labelled with the same symbols (circle and square) for the MDA5^+^ mAbs D**. D)** The predicted selection pressures were obtained for F01 and F12 using the BASELINe computational tool, whereby selection pressures for the complementarity determining region (CDR) and framework region (FWR) (upper left and lower right plots, respectively) and overlays (lower left plot) are visualized for heavy chains. The dotted line depicts the row group, and the solid line represents the column group. The negative selection pressures are shown as green colour and negative values. The intensities of the colours depict the strength of the relative selection pressures. **E)** Dilution curves showing the reactivity of two highly reactive anti-MDA5 clones at different concentration (x-axis) by enzyme-linked immunosorbent assay (ELISA). **F)** The IC50 curves show the in-solution binding of 0.25 μg/mL of mAbs (F01 and F12) to different dilutions of MDA5 antigen (x-axis). **G)** Line blot analysis against different myositis antigens for the two anti-MDA5^+^ mAbs and their corresponding patient serum sample.

Interestingly, based on the index sorting information, the mAb F01 originated from a double negative B cell, while the F12 originated from a switched memory B cell (Figure 2B). Both mAbs originally had IgG1 subclass heavy chains, were VH3 encoded (VH3-7 and VH3-73) and carried lambda light chains. However, they did not share the same clonal origin and there was no similarity in their complementarity determining region 3 (CDR3) amino acid sequences. Strikingly, both mAbs only carried 1-3 predicted somatic hypermutations (SHM) in the heavy and light chain. Although we had expressed some clones (223:01F04, 223:01G11 and 223:01E05) that belong to the same clade as the two MDA5+ mAbs (F01 and F12), none of these mAbs originating from IgM^+^ B cells showed MDA5 positivity in the in-house ELISA when expressed as IgG1 (Figure 2A and C). Using the BASELINe tool to investigate the distribution patterns of replacement/silent mutations as a measurement of selection pressure, both F01 and F12 display negative selection pressure in both the CDR and framework region (FWR) (Figure 2D).

The mAb F01 consistently displayed higher reactivity to MDA5 at lower concentrations as compared to F12 (Figure 2E). Intriguingly, in-solution binding analysis of bivalent IgG showed relatively high apparent affinity of both clones towards recombinant MDA5 with an estimated IC50 of 0.47 nM for F01 and 7.8 nM for F12 (Figure 2F). Importantly, besides binding to the recombinant in-house produced protein, we confirm that both mAbs specifically bind to MDA5 in a commercial line blot assay and showed no binding to other myositis specific or connective tissue disease autoantigens in line blot or antigen microarray assay (Figure 2G, supplementary Figure 3B and C). Moreover, the mAbs showed no cross-reactivity with the functionally related control protein RIG-1 expressed in the same system as the in-house produced MDA5 (supplementary Fig 4).

### MDA5 mAbs recognise distinct epitopes within the Hel2i domain

To characterize the MDA5 mAbs in more detail, we set out to understand the recognised epitopes. Firstly, we performed domain scanning by ELISA and Western blot using different recombinant protein constructs encompassing different domains of the MDA5 protein (Figure 3A). Both F01 and F12 bound to construct A, that represent the full-length MDA5 without the CARD1 domain, as well as constructs D (Hel1-Hel2i-Hel2), E (Hel1-Hel2i-Hel2-pincer) and F (Hel2i domain) (Figure 3B and C, supplementary Figure 4). No binding was seen for the constructs C, G or H that contained only the hel1 domain, the CTD domain or the pincer-CTD region, respectively. Hence, we concluded that the epitopes for both clones are likely located within the Hel2i domain which was the common denominator for binding. While there were some differences in reactivity, both clones bound both in ELISA and Western blot which may suggest targeting of linear epitopes.

**Figure 3.**
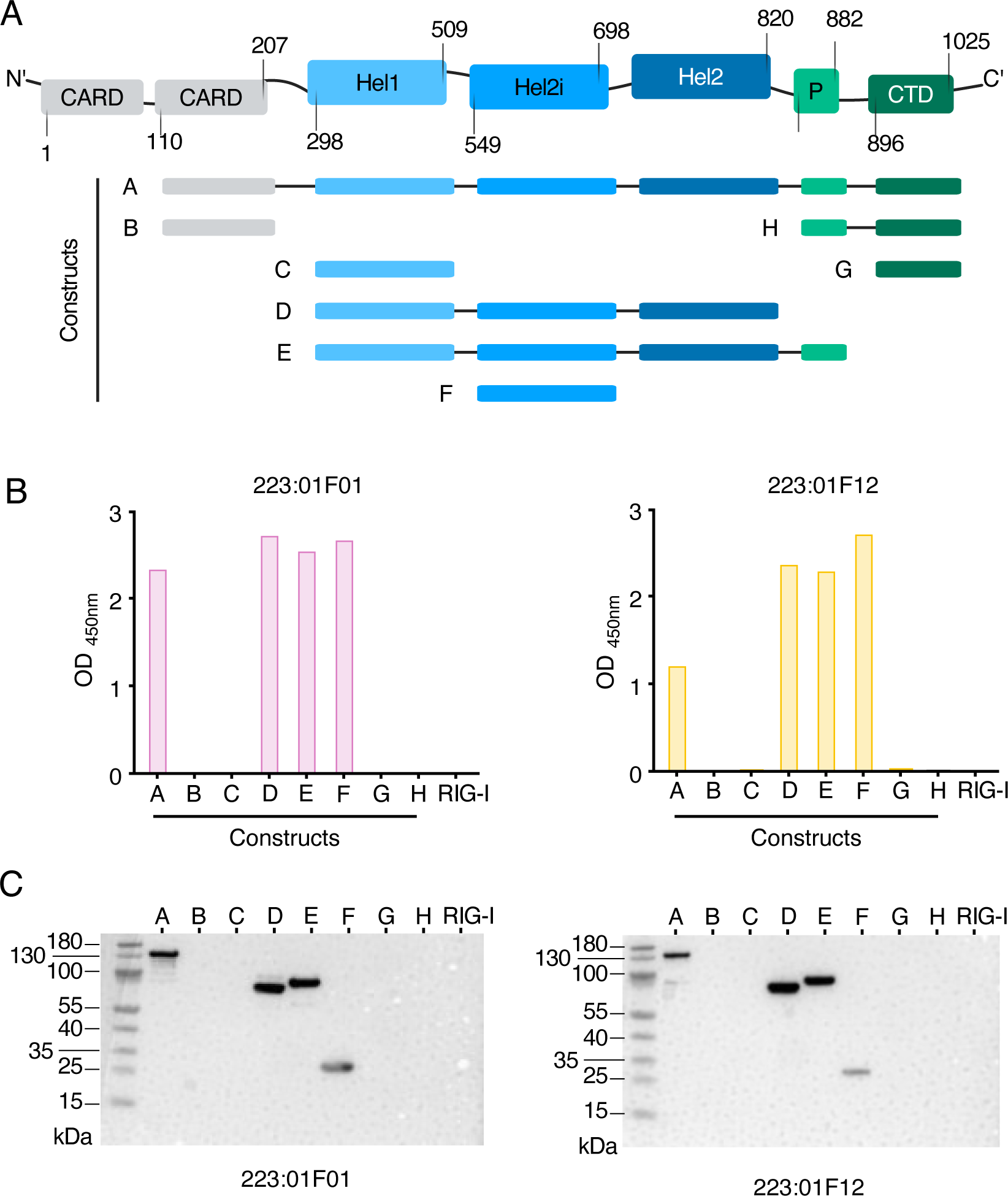
MDA5 domain scanning of mAb binding. **A)** Schematic representation of the MDA5 protein constructs used for ELISAs **B)** The Bar graph shows the ELISA OD for the two mAbs F01 and F12 against the different MDA5 constructs. **C)** Western blot analysis to assess the reactivity of the mAbs against linearized epitopes on the MDA5 constructs A–H and RIG-I

To further characterise the epitopes, we performed peptide scanning with overlapping peptides spanning the MDA5 protein as well as unbiased epitope finger printing using peptide phage display (Figure 4A-D). The peptide scanning highlighted one binding region in Hel2i for the mAb F12 (Figure 4B) which was confirmed and further refined by the epitope fingerprinting (Figure 4D). The AlphaFold structure of MDA5 (acc. no. Q9BYX4) suggests that the epitope is part of a flexible loop (Figure 4C, supplementary Figure 5). Notably, the residues of this loop were not included in the available crystal structure of the protein, further supporting an unstructured region [7]. The binding was confirmed by generating a synthetic peptide covering the epitope (MDA5_645-658_). F12 bound strongly to the peptide while F01 showed no binding (Figure 4E). F12 bound a cyclic version of the peptide only slightly better than a linear variant with cysteine replaced by serine (supplementary Figure 6A), and this position is also not conserved in the results from epitope fingerprinting. Moreover, F12 showed no binding to a control peptide spanning a different MDA5 region (MDA5_659-670_) (Figure 4E).

**Figure 4.**
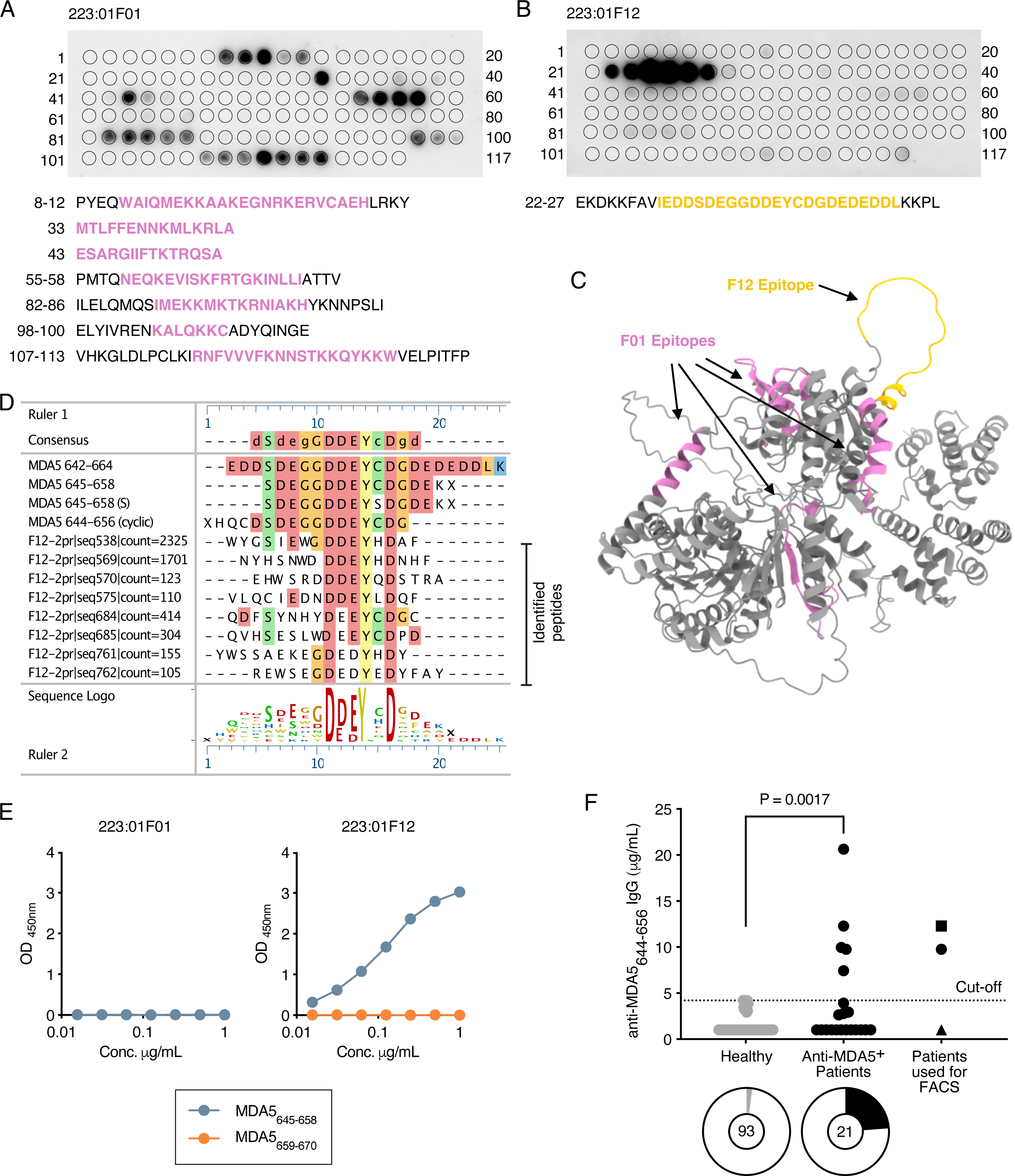
Epitope mapping of anti-MDA5 positive monoclonal antibodies. **A**) Epitope mapping of F01 antibody using overlapping peptides (PepSpot) shows that this antibody recognizes multiple epitopes on the helicase 2i domain. Number on the sides indicate spot numbers. **B**) Epitope mapping of F12 antibody using overlapping peptides (PepSpot) shows that this antibody binds to a linear peptide. **C**) The epitopes bound by the mAb F01 and F12 are highlighted in the MDA5 structure predicted using Alphafold3. **D**) F01 epitope fingerprinting using phage display. **E)** Confirmation of F12 but not F01 to a MDA5_645-658_ synthetic peptide using ELISA. **F**) Serological screening of patient IgG binding to the MDA5_644-656(cycl)_ peptide (n= 93 healthy controls, n= 21 patients).

In contrast to the specificity at the protein level, the binding of the F01 antibody at the peptide level showed multiple bindings to peptides from both within and outside of the Hel2i domain (Figure 4A). There was binding to peptides with positively charged residues, but they shared no consensus motif beyond the charge. Moreover, the phage display epitope fingerprinting did show a similar diversity of enriched motifs. Nevertheless, F01 displayed high apparent affinity for the MDA5 protein, and the binding was independent of potential co-factors such as ions or RNA, as no difference in binding could be detected after addition of EDTA or RNase (supplementary Figure 6B, C and D).

### Serological confirmation of autoantibody recognition of the linear MDA5 peptide epitope in DM

To validate the presence of serum IgG against the novel MDA5-F12 epitope identified from the epitope mapping, we set up an in-house ELISA using a cyclic MDA5_644-656_ peptide for patient serology. Analysing a cohort of 21 anti-MDA5^+^ DM patients, we found that 23.8 % were positive as compared to 2% among healthy individuals (Figure 4F). Of the three patient serum samples that were used for the B cell sorting, two were positive, including the patient from whom the monoclonals were derived. Importantly, the levels of anti-MDA5_644-656_ antibodies were significantly higher in patients compared to healthy controls. These data indicate that antibody response to the MDA5 protein can be directed towards several epitopes and may differ between patients.

## DISCUSSION

This study delineates the phenotype and epitope specificity of MDA5 autoreactive B cells, revealing unusual B cell features combining low somatic hypermutation with high apparent affinity for distinct epitopes within the immunodominant Hel2i domain. Despite central role of autoantibodies in diagnosis of MDA5-DM, origin of autoantibody producing B cells, as well as mechanisms of escaping immune tolerance, are yet to be understood. The recent very encouraging clinical responses with CD19^+^ B cell and B cell maturation antigen (BCMA) plasma cell depleting therapies in autoimmune diseases have further highlighted B cells and antibodies in the disease pathogenesis and call for identification of antigen-specific B cells [24, 25]. We demonstrate that rare MDA5 autoreactive B cells can be detected and isolated from patients with DM using probe enrichment, and that subsequent validation of antigen-reactivity e.g. by BCR cloning and production of recombinant mAbs, is of utmost importance. While prior studies have analysed the presence of serum autoantibodies, in this study using single cell methodology, we produced the first two patient-derived monoclonal anti-MDA5 Abs that recognize the MDA5 helicase domains.

Our results show that two separate B cell lineages from the same patient, reflected by the two MDA5-reactive mAbs encoded by different VH-VL genes, had evolved to primarily target the central helicase domains of the MDA5 protein. Targeting of Hel2i is in-line with previous findings that the helicase domains are the main target of polyclonal anti-MDA5 autoantibodies in patient sera [1, 16, 26].

Using different epitope mapping approaches, clone 223:01F12 showed specific binding to a distinct linear Hel2i peptide epitope. Although, 223:01F01 bound specifically to Hel2i in Western blot and other protein-based tests, the data from peptide arrays would also suggest a conformational epitope with interactions also outside of Hel2i. Intriguing, peptides in several different surface showed binding could suggest separate paratope-epitopes or an epitope less dependent on peptide sequence (e.g. positive charge or motif based interactions). This explains why the phage display approach enriched multiple different peptide sequences with positive charges. Charge dominated interactions with the epitope can easily generate such imprecision at the peptide level. Importantly, the clone showed no unspecific polyreactivity in any control assays and it displayed strong positivity in clinical MDA5 line-blot assay.

Interestingly, both MDA5^+^ mAbs originated from IgG1 BCR sequences and IgG1 have been described as the most common IgG isotype among anti-MDA5 autoantibodies [27]. IgG1 isotype antibodies are potent activators of the complement system [28] and recently an increased amount of the complement component C3 was identified in lung biopsies of patients with MDA5^+^ DM [29]. These findings suggest potential immune complex deposition in the lungs of patients and that the complement system and IgG1 (auto-) antibodies may play a direct or indirect role in tissue specific pathogenesis, evident by pulmonary and dermatological manifestations in anti-MDA5 DM.

The low number of SHM and high apparent affinity of the BCRs suggest that the B cells class-switched without going through (extensive) affinity maturation in the germinal centres, meaning they could have matured through the non-classical extrafollicular B cell maturation pathway [30, 31]. Autoreactive B cells originating from extrafollicular responses have been described in other autoimmune diseases such as lupus nephritis [32]. However, more sequence data from MDA5-reactive BCRs, in-depth B cell phenotyping, and quantitative methods to determine the affinity and avidity are needed to confirm these findings. Several IgM MDA5^+^ B cells were also sorted, cloned and expressed as IgG1 monoclonals. They showed low MDA5 binding; however, this is not surprising as IgM BCRs typically have low affinity and may not retain detectable binding when expressed in the IgG format. Further studies should also address the reactivity of these antibodies.

Since the helicase domains are the enzymatically active sites of the MDA5 protein, and it has been shown for other autoantigens that autoantibodies often bind enzymatically active sites [33–35], we speculate that the antibodies could affect the canonical function of MDA5. This could cause dysregulation of the IFN pathway, which has recently been implicated in the pathogenesis of MDA5^+^ DM in multiple studies [36, 37]. One study showed that B cells from peripheral blood of patients with MDA5^+^ DM had increased type I IFN signalling [36] and another study showed that autoantibodies from MDA5^+^ DM can induce type II IFN in vitro [37]. Whether this is specifically related to anti-MDA5 autoantibodies, dysfunction of canonical MDA5 signalling pathways or other reasons is not known. Moreover, since MDA5 is an intracellular protein during physiological conditions, the question of how and when autoantibodies could access the protein requires further investigation. Additional molecular and in vitro investigations are required to assess this hypothesis. It was recently shown that polyclonal antibodies isolated from some patients with MDA5^+^ DM could affect the canonical functioning of the MDA5 protein, although in this experimental model the underlying mechanisms could not be fully identified [38].

Deconstructing the sequences of BCRs that show high reactivity towards self-antigens (like MDA5) and tracing their lineage would aid understanding of (anti-MDA5) autoantibody pathogenicity and develop antigen-targeted therapies. However, isolating rare autoreactive B cells, especially MDA5-reactive B cells, remains challenging. The efficiency of our strategy was reduced by technical challenges such as the limited availability of biological samples given the rarity of the disease. A recent study found that polyclonal serum antibodies from MDA5+ DM patients recognize multiple linear epitopes of MDA5, with reactivity to specific epitopes correlating with different clinical outcomes [39]. Hence, future studies should investigate differences in epitope recognition at the monoclonal level between very severe and mild ILD cases. The in-depth studied anti-MDA5 clones were both derived from the same patient suffering from interstitial lung disease but not the RP-ILD. Moreover, other clinical disease characteristics, such as lymphopenia may affect the B cell population further decreasing the efficiency of our strategy [40–42]. Of note, at least one of the three patients in this study showed lymphopenia (lymphocytes were below the reference value of 1.1 - 3.5 x10^9^/L, Figure 1A). Although these limitations have slowed the pace for scientific breakthroughs in the past, the knowledge and experience gained throughout this proof-of-concept study will enable the development of more efficient strategies in the future.

In conclusion, we successfully isolated anti-MDA5 autoreactive BCRs and generated the first two MDA5-reactive mAbs, each with distinct antibody characteristics and MDA5-binding characteristics. They can now be harnessed as tools to develop *in vitro* and *in vivo* models to understand the potential pathogenicity of the anti-MDA5 autoantibodies and will help lay the basis for future studies on the MDA5-reactive Adaptive Immune Receptor Repertoire (AIRR) and the development of MDA5-targeted therapies.

## AUTHORS CONTRIBUTIONS

EVG, VM, IEL, CG, KC, VJ were responsible for conceptualization of the study. DD, IEL recruited patients and provided the clinical data. EVG, MG, ND, KK, RS, MH, AVV, EW, SG, BH, VJ performed the experiments. EVG, MG, MS, VM, IEL, BH, CG, KC, VJ were involved in the interpretation of data. EVG, IEL, CG, KC, VJ drafted the initial version of the Manuscript. All authors reviewed, edited, read, and approved the final version of the manuscript.

## ACKNOWLEDGEMENTS

We thank all patients with IIM that contributed to these research efforts. We also wish to thank the research nurses Helene Sandlund and Ingrid Gerhardsson, as well as Gloria Rostvall, Lucymary Okechukwu and Julia Norkko, for managing the cohort biobanking and handling of blood samples. We thank Linda Mathsson-Alm at Thermo Fisher Scientific, Uppsala, Sweden, for the antigen-array analysis of monoclonal antibodies.

