## Supplemental Figures for "Anti-MDA5 monoclonal antibodies from patients with dermatomyositis - B cell characteristics and differential targeting of the helicase domains"

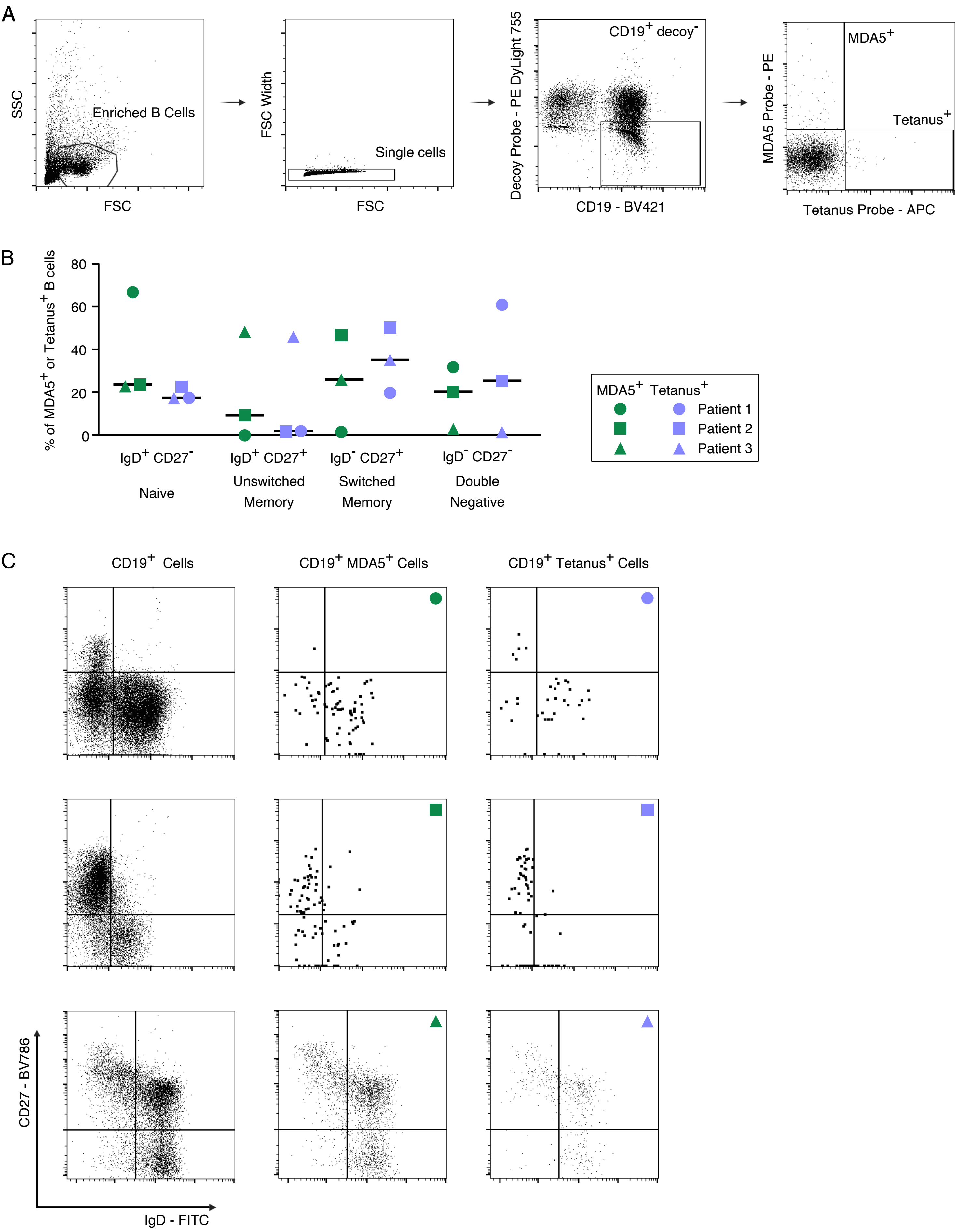


**Supplementary Figure 1: Detection, isolation and phenotyping of anti-MDA+ autoreactive B cells. A)** Flow cytometry gating strategy for detection, phenotyping and sorting of antigen specific $CD19^+$ B cells from PMBC of patients with anti-$MDA5^+$ DM. **B)** B cell sub phenotyping based on surface expression of IgD and CD27 **C)** Flow cytometry dot plot showing surface expression of IgD and CD27 in all $CD19^+$, $CD19^+$ $MDA5^+$ and $CD19^+$ $tetanus^+$ B cells from the 3 patients described in figure 1A.

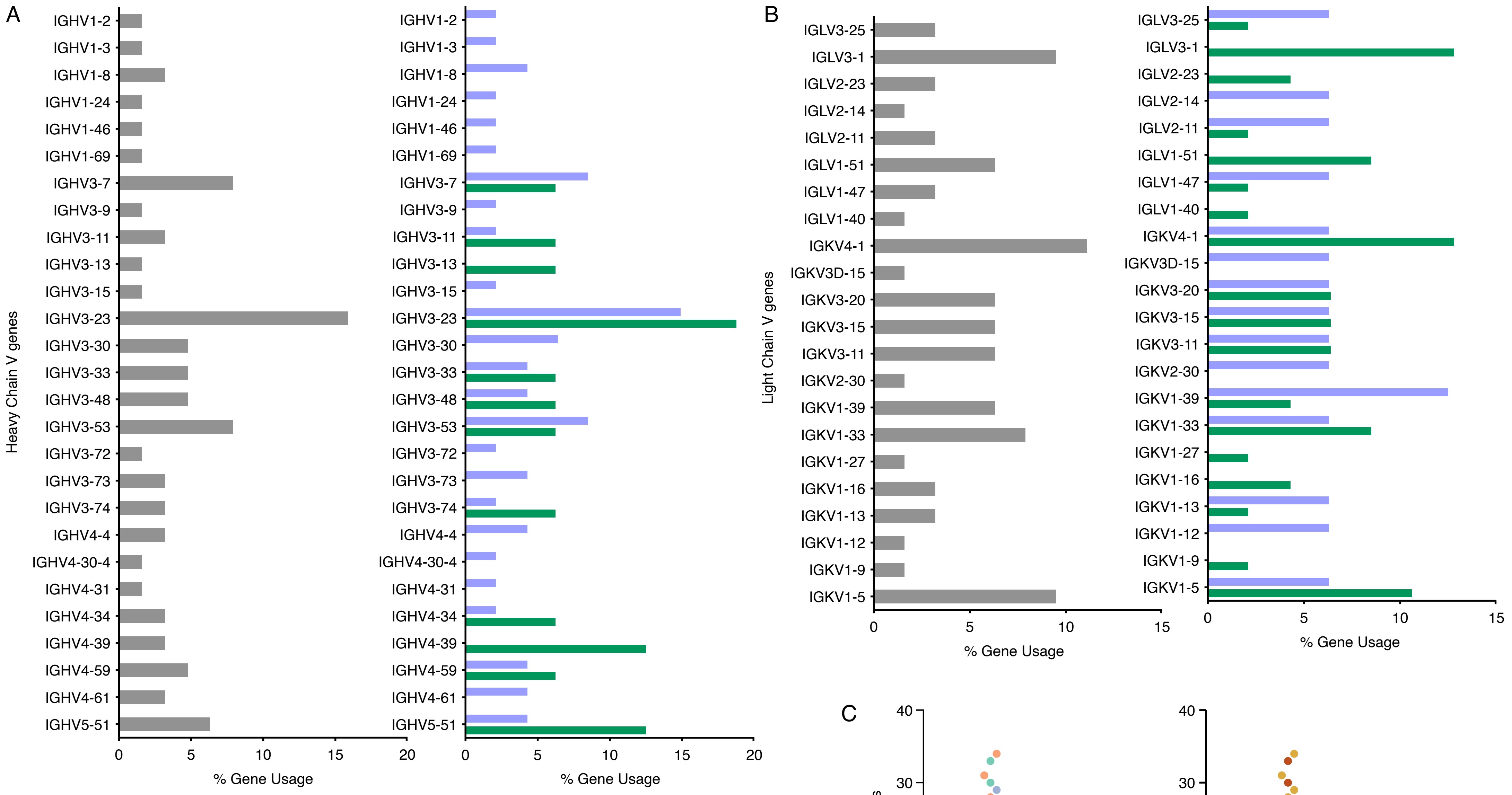


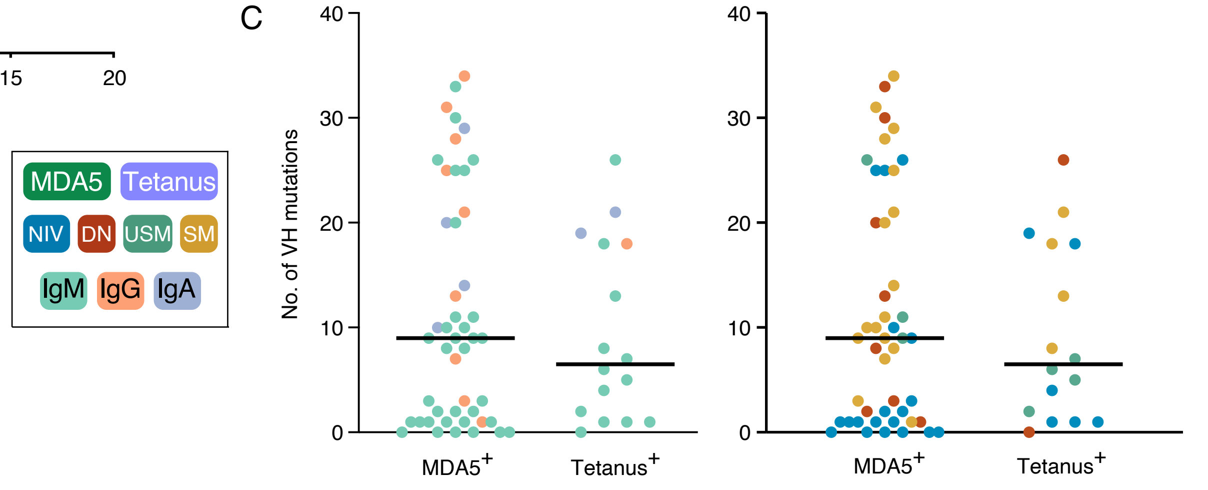


**Supplementary Figure 2: Immunoglobulin repertoires of CD19+ MDA5+ and CD19+ tetanus+ B cells. A)** Left panel shows heavy chain V gene usage of all antigen specific (MDA5+ or tetanus+) CD19+ B cells and right panel is the gene usage stratified on MDA5+ and tetanus+ B cells. B) Light chain V gene usage of all antigen specific (MDA5+ or tetanus+) CD19+ B cells. **C)** Comparison of the number of mutations in the heavy chain V gene between CD19+ MDA5+ and CD19+ tetanus+ B cells. (NIV: Naïve, DN: Double Negative, USM: Unswitched Memory or SM: Switched Memory)

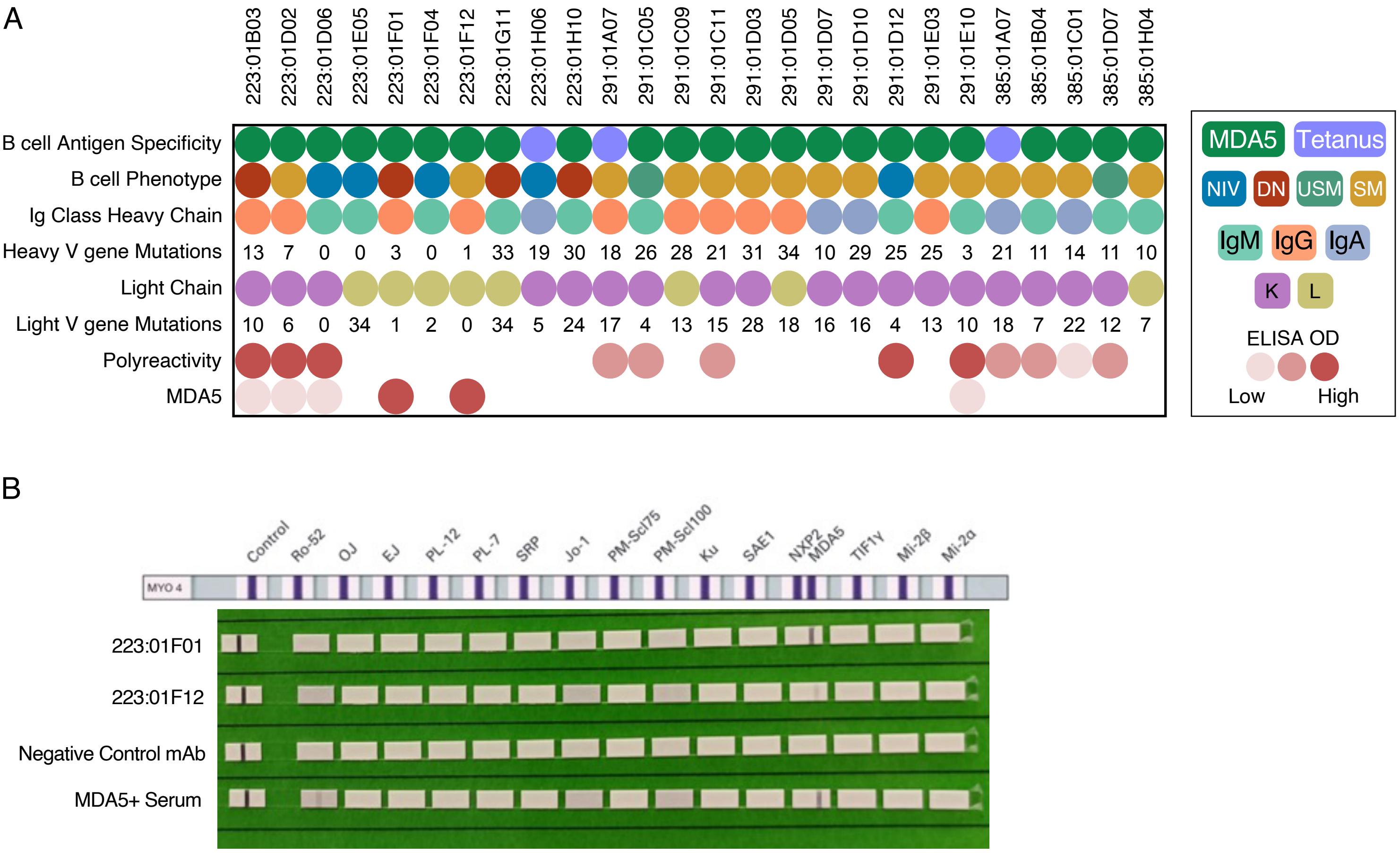


C

| | CENP-B | dsDNA | Fibrillarin | Jo-1 | PCNA | PM/Scl-100 | RIP P0 | RIP P1 | RIP P2 | RNA Pol III | RNP70 | RNP-A | RNP-C | Ro52 | Ro60 | Scl-70 | Sm | SmBB | SS-B/La |
|---|---|---|---|---|---|---|---|---|---|---|---|---|---|---|---|---|---|---|---|
| 223:01F01 | 37 | 37 | 84 | 108 | 153 | 65 | 101 | 36 | 30 | 112 | 52 | 58 | 55 | 54 | 44 | 95 | 43 | 41 | 139 |
| 223:01F12 | 52 | 64 | 107 | 94 | 102 | 73 | 102 | 41 | 33 | 115 | 89 | 65 | 61 | 56 | 52 | 119 | 43 | 54 | 135 |
| ANA+ Serum | 4674 | 9965 | 696 | 5143 | 348 | 1706 | 496 | 472 | 345 | 422 | 4141 | 644 | 3400 | 14663 | 1938 | 1548 | 9299 | 9525 | 1411 |
| Assay Diluent | 43 | 33 | 65 | 152 | 215 | 88 | 170 | 38 | 32 | 162 | 54 | 69 | 78 | 73 | 67 | 114 | 49 | 65 | 257 |

FI: Low – High

Fluorescent Intensity

**Supplementary Figure 3: Characteristics of patient-derived monoclonal antibodies to MDA5. A)** The heatmap shows the characteristics of the 26 BCRs that were expressed as mAbs. The map shows the antigen specificity of the B cells (MDA5$^+$ or tetanus$^+$), B cell phenotype (NIV: Naïve, DN: Double Negative, USM: Unswitched Memory or SM: Switched Memory), Ig isotype (M, G or A) of the BCR, number of heavy chain V gene mutations, light chain type (K:kappa or L:lambda), number of light chain V gene mutations, polyreactivity and MDA5 reactivity based on ELISA. **B)** Line blot image using Euroimmune MYO4 assay containing different myositis antigens for the two anti-MDA5+ mAbs, their corresponding patient serum sample and a negative control mAb produced using the same method. **C)** Heatmap with the fluorescent intensities (FI) values of for the mAb reactivity against different autoantigens.

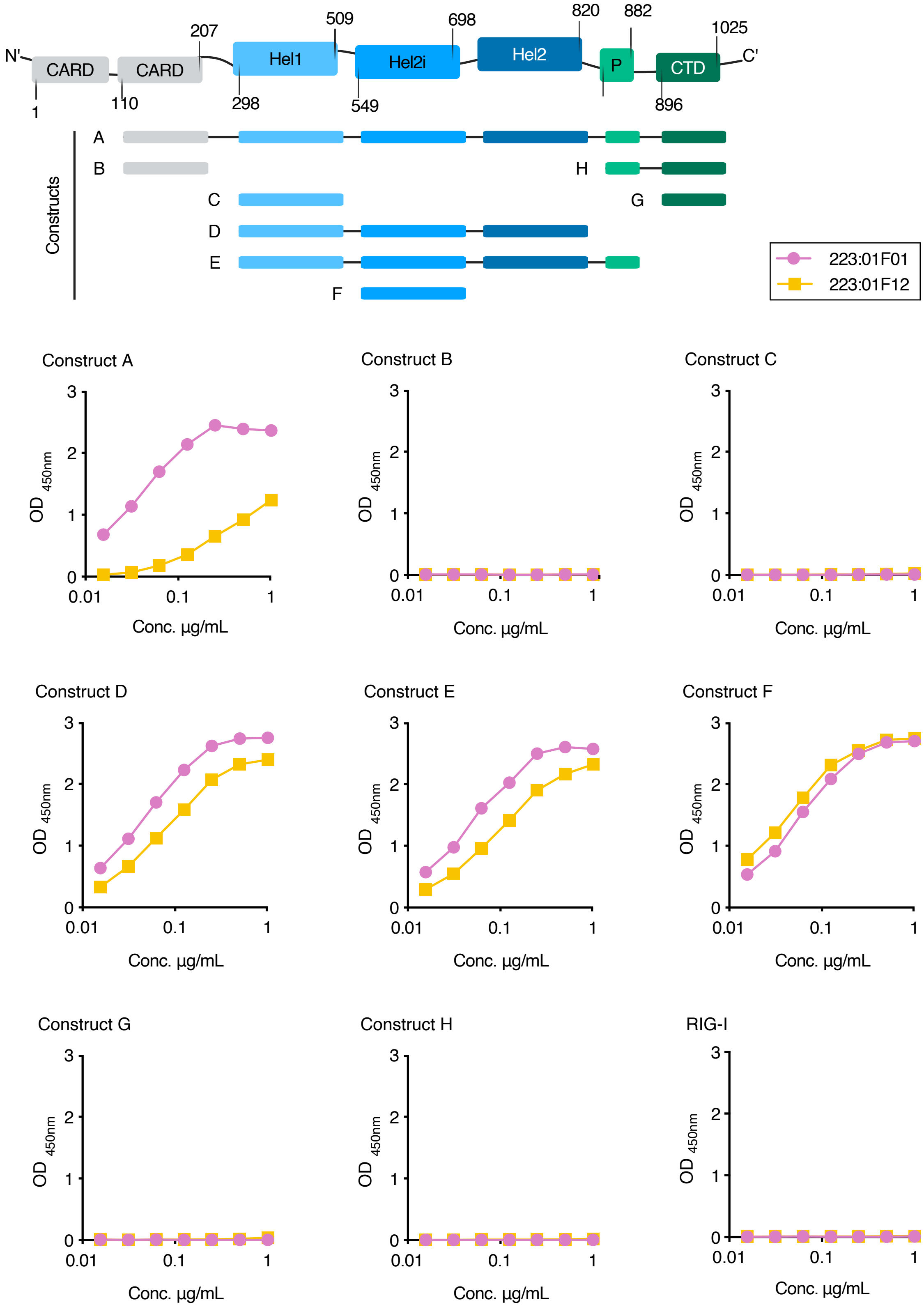


**Supplementary Figure 4: MDA5 domain scanning of mAb binding.** Schematic figure showing MDA5 domains and constructs spanning MDA5 sequence. The plots show the ELISA reactivity of anti-MDA5$^{+}$ monoclonal antibodies 223:01F01 and 223:01F12, at concentration ranging from 1 µg/mL – 0.016 µg/mL towards the different constructs spanning MDA5 and full-length RIG-I.

A

223:01F01

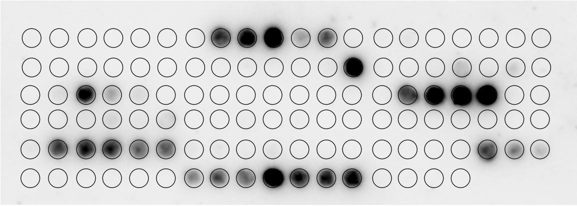


PYEQWAIQ**MEKKAAKEGNRKERV**CAEHLRKY

8 PYEQWAIQMEKKAAK
9 WAIQMEKKAAKEGNR
10 MEKKAAKEGNRKERV
11 AAKEGNRKERVCAEH
12 GNRKERVCAEHLRKY

**MTLFFENNKMLKRLA**

33 MTLFFENNKMLKRLA

**ESARGIIFTKTRQSA**

43 ESARGIIFTKTRQSA

PMTQNEQKEVI**SKFRTGKI**NLLIATTV

55 PMTQNEQKEVISKFR
56 NEQKEVISKFRTGKI
57 EVISKFRTGKINLLI
58 KFRTGKINLLIATTV

ILELQMQS**IMEKKMKTKRNIA**KHYKNNPSLI

82 ILELQMQSIMEKKMK
83 QMQSIMEKKMKTKRN
84 IMEKKMKTKRNIAKH
85 KMKTKRNIAKHYKNN
86 KRNIAKHYKNNPSLI

ELYIVREN**KALQKKC**ADYQINGE

98 ELYIVRENKALQKKC
99 VRENKALQKKCADYQ
100 KALQKKCADYQINGE

VHKGLDLPCLKI**RNFVVVFKNNSTKKQYKKW**VELPITFP

107 VHKGLDLPCLKIRNF
108 LDLPCLKIRNFVVVF
109 CLKIRNFVVVFKNNS
110 RNFVVVFKNNSTKKQ
111 VVFKNNSTKKQYKKW
113 NNSTKKQYKKWVELP
113 KKQYKKWVELPITFP

B

223:01F12

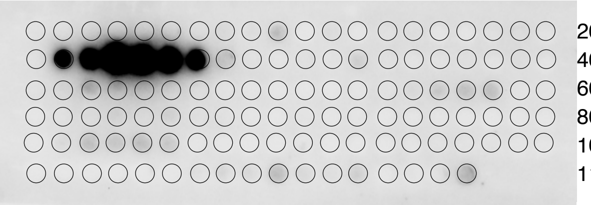


EKDKKFAV**IEDDSDEGGDDEYCDGDEDEDDL**KKPL

22 EKDKKFAVIEDDSDE
23 KFAVIEDDSDEGGDD
24 IEDDSDEGGDDEYCD
25 SDEGGDDEYCDGDED
26 GDDEYCDGDEDEDDL
27 YCDGDEDEDDLKKPL

C

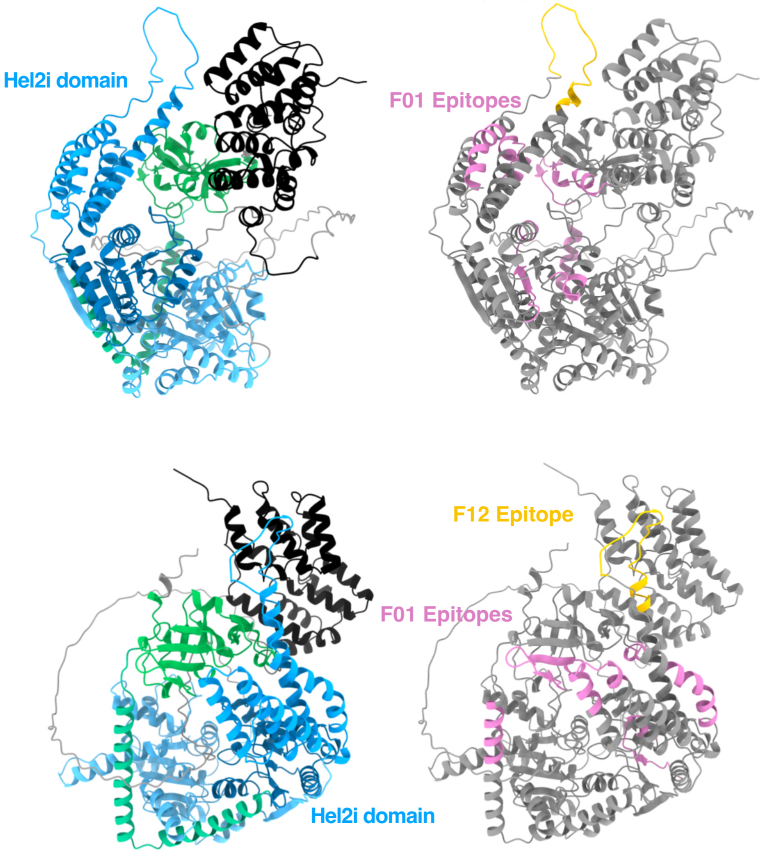


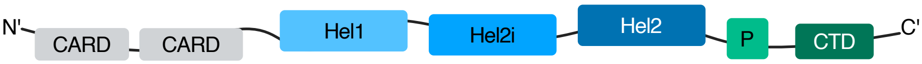


**Supplementary Figure 5: Epitope mapping of anti-MDA5 positive monoclonal antibodies.**

**A-B)** Epitope mapping of anti-MDA5$^{+}$ monoclonal antibodies using PepSpot technology. Chemiluminescent signal images and overlapping peptides' amino acid sequences with designated minimal epitope for 223:01F01 **(A)** and 223:01F12 **(B)**. **C)** AlphaFold 3 prediction structure of MDA5 (acc no AF-Q9BYX4-F1-v4) showing determined F12 and F1 epitopes together with their positions within MDA5 domains.

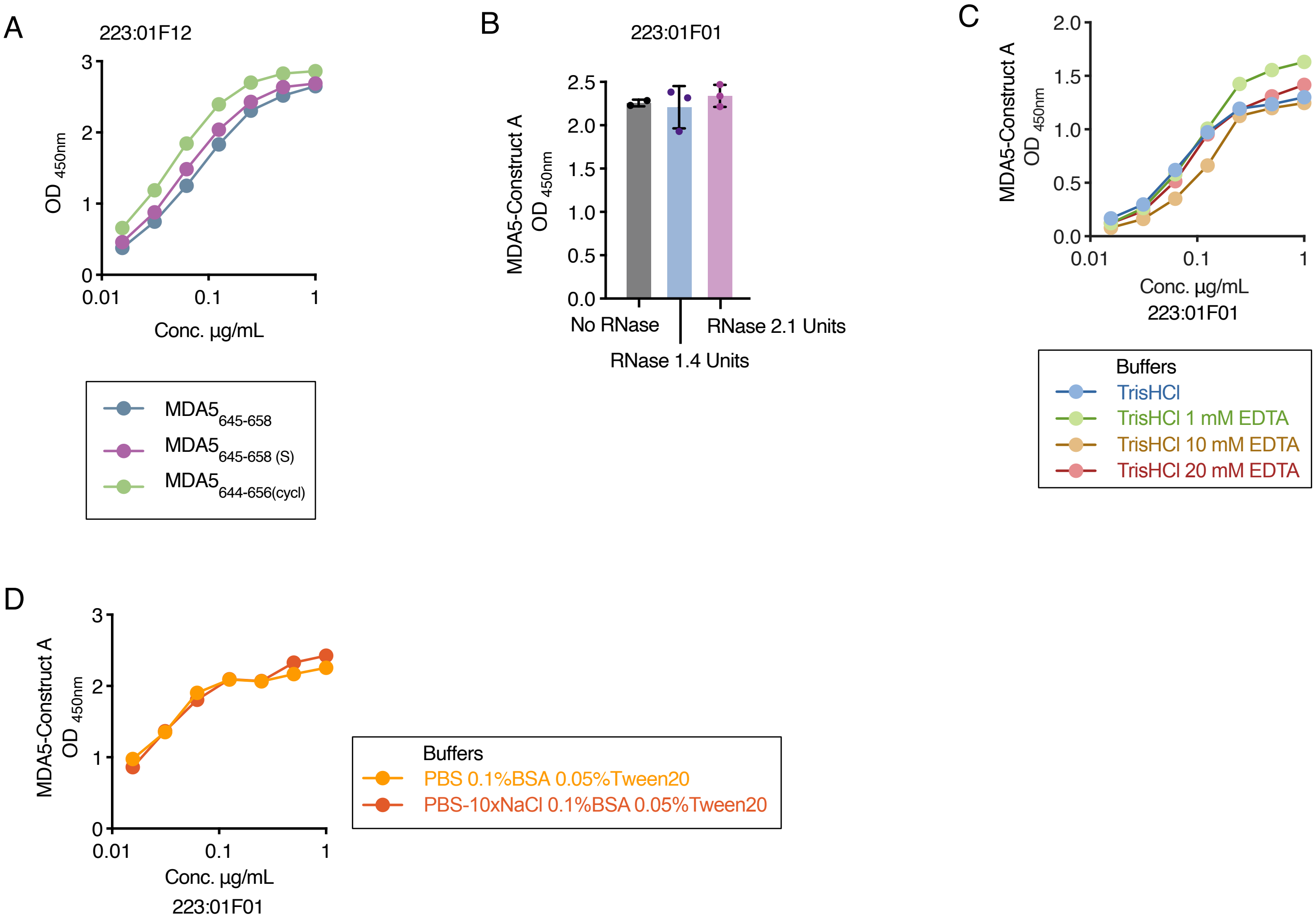


**Supplementary Figure 6: Binding properties of anti-MDA5$^{+}$ mAbs. A)** 223:01F12 binding to synthetic peptide covering the epitope MDA5$_{645\text{-}658}$, cysteine to serine substitution peptide version MDA5$_{645\text{-}658(S)}$, and cyclic version of the peptide MDA5$_{644\text{-}656(cycl)}$ in ELISA. **B)** 223:01F01 mAb reactivity to MDA5-construct A with or without RNase A at mAb concentration of 1 µg/mL. **C-D)** 223:01F01 mAb reactivity to MDA5 construct A in different buffer conditions.

**Supplementary Table 1: MDA5 Constructs used for the ELISA**

Molecular weight and amino acid coverage of MDA5 constructs and RIG-I control protein construct utilized for antibody purification and in ELISA.

| Construct | Domains | MW (kDa) | StartStop |
|---|---|---|---|
| A | CARD(2)-Hel1-Hel2i-Hel2-Pincer-CTD | 109.3 | A110-D1025 |
| B | CARD(2) | 16.2 | A110-A207 |
| C | Hel1 | 38.3 | A298-L509 |
| D | Hel1-Hel2i-Hel2 | 74.4 | A298-R820 |
| E | Hel1-Hel2i-Hel2-Pincer | 81.5 | A298-M882 |
| F | Hel2i | 36.8 | P549-N698 |
| G | CTD | 20.2 | Y896-D1025 |
| H | Pincer-CTD | 29.5 | M816-D1025 |
| RIG-I | RIG-I | 84.7 | S230-K925 |

MW, molecular weight; CARD, caspase recruitment domain; Hel, Helicase Domain; CTD, C-terminal domain, RIG-I, retinoic acid-inducible gene I.

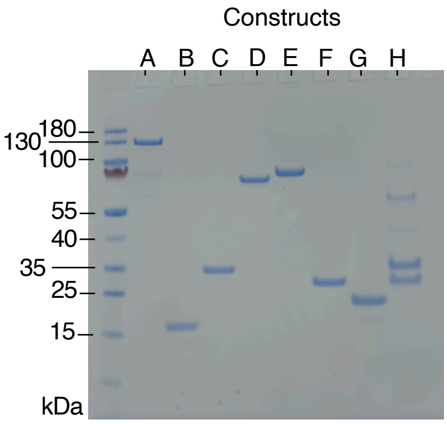


SDS-PAGE gel showing the size shift of the different constructs

**Supplementary Table 2: Antibodies used for FACS**

| Marker | Conjugate | Clone | Manufacturer |
|---|---|---|---|
| CD3 | APC-H7 | SK7 | BD Pharmingen™ |
| CD14 | APC-H7 | MϕP9 | BD Pharmingen™ |
| CD16 | APC-H7 | 3G8 | BD Pharmingen™ |
| CD19 | BV421 | HIB19 | BD Horizon™ |
| CD27 | BV786 | L128 | BD Horizon™ |
| IgD | FITC | IA6-2 | BD Pharmingen™ |
| CD38 | BV711 | HIT2 | BD Horizon™ |
